# Human iPSC-Vascular Smooth Muscle Cell Spheroids Demonstrate Size-dependent Alterations in Cellular Viability and Secretory Function

**DOI:** 10.1101/2022.03.31.486610

**Authors:** Sara Islam, Jackson Parker, Biraja C. Dash, Henry Hsia

## Abstract

Human induced pluripotent stem cells and their differentiated vascular cells have been revolutionizing the field of regenerative wound healing. These cells are shown to be rejuvenated with immense potentials in secreting paracrine factors. Recently, hiPSC-derived vascular smooth muscle cells (hiPSC-VSMC) have shown regenerative wound healing ability via their paracrine secretion. The quest to modulate the secretory function of these hiPSC-VSMC is an ongoing effort and involves the use of both biochemical and biophysical stimuli. This study explores the development and optimization of a reproducible, inexpensive protocol to form hiPSC-VSMC derived spheroids to investigate the implications of spheroid size on viability and paracrine secretion. Our data shows the successful formation of different sizes of spheroids using various amount of hiPSC-VSMC. The hiPSC-VSMC spheroids formed with 10000 cells strike an ideal balance between overall cell health and maximal paracrine secretion. The conditioned medium from these spheroids was found to be bioactive in enhancing human dermal fibroblast cell proliferation and migration. This research will inform future studies on the optimal spheroid size for regenerative wound healing applications.

## Introduction

Human induced-pluripotent stem cells (hiPSCs) have been key to advancements in the fields of bioengineering and tissue regeneration. hiPSCs provide researchers with an unlimited, pure supply of cells with enormous regenerative potential and without the ethical concerns traditionally associated with embryonic stem cells.^1^ They have been used extensively in cell therapy, disease modeling, and drug discovery.^2^ However, a major obstacle to clinical use of these cells is their reduced therapeutic potential and rapid death following transplantion.^3-7^ Other restrictions to direct cell use have been limits in cell expansion, migration from the targeted therapy site, and inability to withstand hostile microenvironments.^8,9^ Therefore, there is mounting interest in improving the survival and efficacy of stem cell therapies *in vivo*.^7^

In recent years, research has shown that cell-mediated regeneration is a result of the cell’s secretome. The cell secretome contains growth factors, cytokines, extracellular vesicles, and exosomes which guide and support cellular function.^7,10^ Use of the hiPSC-derived secretome in lieu of cells has allowed researchers to overcome obstacles relating to in-culture cell expansion and post-transplantation cell death.^3,4,6^ Additionally, the hiPSC-derived secretome is cell-free and carries less risk of immune reactivity.^11^ Previous studies have demonstrated that the hiPSC-derived secretome can be used to treat hematopoietic disease, heart failure, liver failure, bone defects, and cartilage defects.^7^ With all of these applications in mind, researchers are constantly looking for ways to increase the production of paracrine secretion and in turn, the therapeutic potential of hiPSCs.

To this end, many studies have found that culturing cells in a 3D spheroid environment enhances their therapeutic potential and survival.^3^ These 3D spheroids are self-assembled aggregates of cells and more closely resemble the tissue microenvironment. To date, several different stem cell lineages have been formed into spheroids including hiPSCs and mesenchymal stem cells (MSCs) derived from umbilical cord tissue, endothelial cells, hepatocytes, adipose, smooth muscle, bone marrow, and menstrual blood.^5,8,10-15^ 3D spheroids have been used to promote cartilage regeneration, treat lung injury, and model epithelial cell behavior.^13,16,17^ In comparison to the traditional culture technique of cells in a monolayer, cells cultured as spheroids have demonstrated enhanced cell-survival, paracrine factor secretion, and tissue forming potential.^3,4,6,8,9,18-21^ They also have higher levels of tissue-specific gene expression and increased exosome production.^10,22^ Overall, spheroids enhance biological processes relating to immunity, inflammation, angiogenesis, and wound healing. ^14,23-26^ A previous study by Murphy *et. al*. reported that spheroid secretome increased the function of macrophages and endothelial cells and even accelerated wound healing in an *in vitro* skin model.^25^ Being able to increase production via spheroid formation and then utilize the secretome for management of chronic wounds is an exciting prospect.

The basic mechanisms contributing to spheroid formation can be found in many naturally occurring settings including embryogenesis, organogenesis, and tumor formation.^27,28^ Spheroid assembly occurs without the need for extra stimuli, but is heavily reliant on cadherin and integrin expression.^8,29,30^ Studies have even shown that it is the interaction of integrin with the ECM that determines the rate of spheroid formation.^31^ Since this is a cell-mediated self-assembly process, there is no one way to induce spheroid formation and several methods exist. The oldest, and most traditional technique has been the hanging drop method.^21^ Other variations to spheroid creation include low attachment microplates, liquid overlay, spinner flask, and forced-aggregation techniques.^21,31-33^ Developments in microfluidics have been the most recent advancement and allow for more precise spheroid fabrication, but have the major drawback of requiring costly specialized equipment.^27,34,35^

Our previous work has shown that the hiPSC-VSMC secretome improves cell-mediated regenerative healing.^1,36-38^ We showed that their secretome improved wound closure and angiogenesis via secretion of proangiogenic growth factors in both acute and diabetic wound models.^36-40^ Others have used hiPSC-VSMC to develop structures resembling microvasculature to be used in tissue engineering.^41,42^ The benefits of hiPSC-VSMC encourages us to explore mechanisms to modulate the paracrine secretion and in turn, their therapeutic potential in wound healing.

The overall goal of this study was to culture hiPSC-VSMC as 3D spheroids to maximize their viability and secretome production. The objectives were to i) develop an inexpensive and reproducible protocol to form hiPSC-VSMC spheroids and ii) investigate the optimal size of hiPSC-VSMC spheroids for regenerative wound healing applications. For our experiments, we used agarose-coated, u-bottom microwells for the hiPSC-VSMC of various amounts to self-aggregate to spheroids of varying sizes. Our protocol does not require centrifugation, and thus avoids the potential damage to cells through shearing forces. We found our spheroids maintained the hiPSC-VSMC phenotype across all sizes. Most importantly, we optimized the 10000-cell size of spheroids to be the size which balances both viability and paracrine secretion.

## Methods

### hiPSC-VSMC Differentiation and Characterization

A pure population of hiPSC-VSMC was generated using a previously established protocol.^1,36-38^ In brief, hiPSCs were cultured in feeder-free conditions until reaching 80% confluency. At this point, embryoid bodies (EB) were made on 6-well, ultra-low attachment plates and cultured for 5 days with EB differentiation medium. After another 5 days on gelatin-coated plates, attached cells were transferred to Matrigel-coated plates and cultured with smooth muscle growth medium (SmGM-2) until 80% confluency.

hiPSC-VSMC were grown on 0.1% gelatin-coated plates using SmGM-2 medium supplemented with 1% antibiotic-antimycotic. Growth medium was changed every other day until 80% confluency was reached. The differentiated cells were characterized using immunofluorescence staining for major smooth muscle cell markers calponin, smooth muscle alpha-actin (SMA), and SM-22α. A passage of no more than 10 was used for all experiments.

### Spheroid Formation

Self-aggregating, hiPSC-VSMC spheroids were produced in the following steps (Figure 1A). A 96-well, U-bottom plate was coated with 50 µL of 2% w/v of liquid agarose and allowed to solidify for 30 minutes at room temperature. 200 µL of SmGM-2 medium was then added to each of the wells. Cells of final amounts 5000, 10000, 20000, and 40000 cells where then added to form spheroids. The plate was allowed to remain in the incubator undisturbed for three days, with cells aggregating into spheroids after 24 hours. On day 3, conditioned medium (CM) was collected and spheroids were fixed with 4% paraofrmaldehyde (PFA) for at least 15 minutes in preparation for immunostaining.

**Figure 1:**
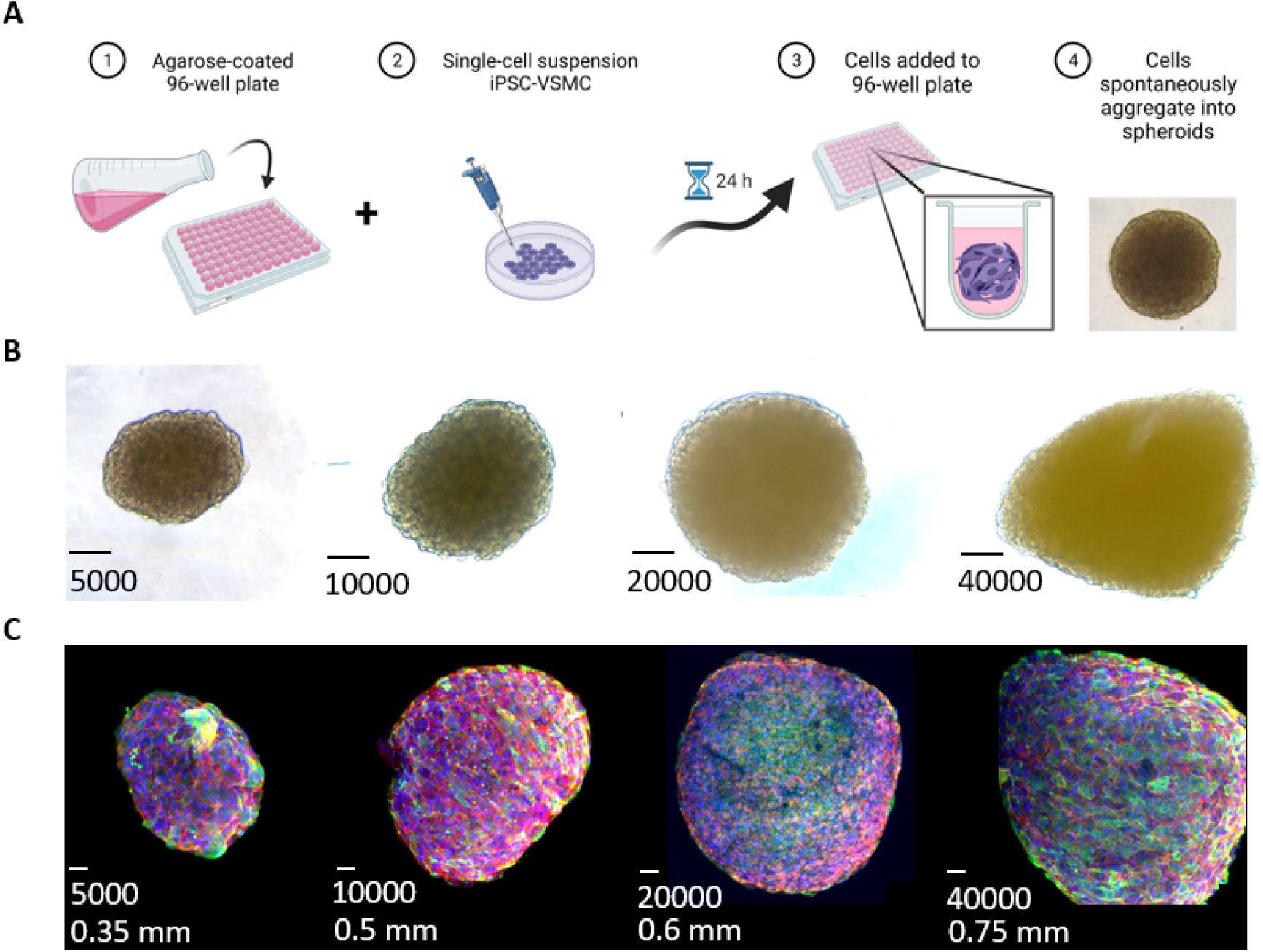
Spheroid formation. (A) Schematic demonstrating the process of spheroid formation (made with BioRender). (B) Representative bright field images of different size spheroids made using 5000, 10000, 20000 and 40000 hiPSC-VSMCs. Scale bar measures 50 μm. (C) Immunofluorescence images of the four sizes of spheroids used for all experimental assays with their Immunofluorescence images stained with Calponin (green), rhodamine phalloidin stains cytoskeleton (red), and Dapi stains nuclei (blue). Scale bar measures 50 μm.

### AlamarBlue Assay

Spheroids of varying sizes (5000, 10000, 20000, and 40000 cells) were tested for cell viability 72 hours after formation using an AlamarBlue assay. Spheroids were incubated with 20 µL of AlamarBlue reagent (ThermoFisher) added directly to their respective wells containing 200 µL of SmGM-2 medium. The plate was incubated in the cell culture incubator for 2 hours at 37LJ. Results were determined using a plate reader with fluorescence intensity at 540 nm excitation and 590 emission wavelengths. Cell viability was normalized using the average fluorescence intensity value of the 5000-cell spheroids.

### Live/Dead Assay

The live/dead assay was performed using the LIVE/DEAD® Cell Imaging Kit according to the manufacturer’s protocol.^1,36-38^ 72 hours post-creation, 0.2 µL of ethidium homodimer-1 (EthD1) dead red was added directly to the 96-well plate containing spheroids of 5000, 10000, 20000, and 40000 cells. The samples were then incubated for 30 minutes at room temperature. After incubation, spheroids were washed 3x with PBST (PBS + 0.05% Tween-20) and fixed with 4% PFA for at least 15 minutes. The samples were counterstained with dapi before imaging on a confocal microscope. The spheroids were ultimately too dense to count the number of live cells stained with live green. Thus, the percentage of dead cells was calculated by counting the number of cells positively stained with EthD1 versus the number of nuclei in 5 different frames of reference for each spheroid.

### Cell Proliferation Assay

The cell proliferation assay was conducted according to the manufacturer’s protocol.^43^ On day 1 of spheroid formation, 1 µL of 10 mM BrdU labeling solution was added directly to the wells containing 200 µL of SmGM-2 medium and spheroids with the following sizes: 5000, 10000, 20000, and 40000 cells. On day three, after spheroids were collected and fixed with 4% PFA, samples were blocked with 2.5% w/v bovine serum albumin for 1 hour. Then, anti-BrdU primary antibody was added at a concentration of 1:100 and left to incubate at 4LJ overnight. The following day, samples were washed 3x with PBST before being incubated with Alexa Fluor 488 secondary antibody for 1 hour at room temperature. The cytoskeleton was stained with rhodamine phalloidin, and nuclei were counterstained with dapi before imaging on a confocal microscope. The percentage of proliferating cells was determined by counting the number of cells positive for BrdU and comparing this to the total number of nuclei in 5 different frames of reference for each sample.

### Lactate dehydrogenase assay

Lactate dehydrogenase (LDH) cytotoxicity assay was done using the CyQUANT™ LDH Cytotoxicity Assay and performed according to the manufacturer’s protocol.^36-38^ Conditioned medium from the 3-day culture 5000, 10000, 20000, and 40000 cell samples was mixed with 50 µL of LDH substrate mix and incubated at 20LJ for 30 minutes. The LDH levels in each sample were quantified using a plate reader set to 490 nm absorbance. Cell cytotoxicity was normalized using the average absorbance value of the 5000-cell spheroids.

### Immunofluorescence Staining

hiPSC-VSMC spheroids harvested and fixed with 4% PFA were further processed for phenotypic characterization by immunofluorescence staining. Spheroids were first blocked with 100 µL of 2.5% BSA for 1 hour at room temperature. The primary antibodies targeted the smooth muscle cell markers calponin, smooth muscle α-actin (SMA), and SM-22α. Primary antibody was added at a dilution of 1:100 and samples were incubated at 4LJ overnight. The next day, spheroids were washed three times with PBST. Secondary antibody (1:250) tagged with Alexa Fluor 488 and Alexa Fluor 555 was then added to the spheroids. These were allowed to incubate on a plate shaker for 1 hour at room temperature. Dapi was used as a counterstain and rhodamine phalloidin was used to stain the cytoskeleton. The spheroids were washed a final three times with PBST before mounting the samples on to slides for confocal imaging.

### ELISA

Conditioned media (CM) was collected from the spheroids 72 hours post-creation and used to characterize their secretory profiles in terms of various cytokines, growth factors, and proteases. The factors tested were as follows: vascular endothelial growth factor (VEGF), basic fibroblast growth factor (bFGF), angiopoietin (Ang)-1, transforming growth factor (TGF)-β, tumor necrosis factor (TNF)-α, interleukin (IL)-10, stromal cell-derived growth factor (SDF)-1α, and matrix metalloproteinase (MMP)-2. 85 µL of CM was added to microwells on a 96-well ELISA plate in duplicate and allowed to incubate overnight at 4LJ. The following day, CM was removed, and the samples were incubated with 100 µL of 2.5% BSA for 1 hour at room temperature. Samples were subsequently washed with PBST three times before the addition of primary antibody (1:2500) and once again placed in the fridge to incubate overnight. Samples were then once again washed three times with PBST and then incubated with secondary antibody (1:2500) for two hours at room temperature. The plates were washed another three times with PBST and 100 µL of TMB substrate solution was added to the plates. The TMB substrate was incubated on a plate shaker at room temperature for 20 minutes. Then, 100 µL of 2N HCl stop solution was promptly added to each microwell. Results were measured at an absorbance of 450 nm using a plate reader. ELISA data was normalized using the average absorbance value of the 5000-cell spheroids.

### Fibroblast Proliferation, Cytotoxicity, and Paracrine Secretion

Primary human dermal skin fibroblasts were grown in fibroblast media (DMEM high glucose + 5% FBS + 1% antibiotic-antimycotic) until 80% confluency. The cells were then dissociated. 50000 cells were transferred into each well of a 24-well plate for functional assays. Fibroblasts were treated with three different types of medium. Treatment with basal DMEM high glucose (DMEM) was used as the negative control and fibroblast conditioned medium (Fibroblast CM) was used as the positive control. CM from 10000-cell hiPSC-VSMC spheroids (Spheroid CM) served as the experimental group. The cells were allowed to incubate in their respective medium at 37LJ for 24 hours.

Proliferation was assessed using an AlamarBlue assay. Briefly, media was removed from the samples and set aside for ELISA and LDH experiments. 200 µL of AlamarBlue reagent in DMEM was added to each microwell. The plate was returned to the cell incubator for 2 hours and results were determined as described earlier. Cell proliferation was normalized using the average fluorescence intensity value of the DMEM negative control group.

Reserved media was used for a LDH cytotoxicity assay and ELISA as described earlier. We evaluated paracrine secretion of bFGF and MMP-2.

### Migration Assay

For the migration assay, 50000 primary human dermal fibroblasts were seeded into each well of a 24-well plate. After day 1 of culture, a 1000 µL pipette tip was used to scratch one line down the center of each well. Bright field images of each sample were taken as a reference to before migration. Next, fibroblasts were treated with three different types of media: DMEM, Fibroblast, and Spheroid CM. The plate was then allowed to incubate for 24 hours at 37LJ. The following day, bright field images were taken to capture the cell migration of each treatment group. Relative migration was quantified by counting the number of cells that entered the scratch zone when comparing bright field images from before and after incubation with treatment media.

### Statistical Analysis

Statistical analysis was done using the Graph Pad Prism 9 software. Significance was determined using one-way ANOVA and student *t*-test. For all experiments, the *n* number was at least 4-27 and the data was presented using the mean +/-the standard deviation.

## Results

### hiPSC-VSMC spheroids retain their smooth muscle phenotype and morphology

A single-cell suspension of hiPSC-VSMC, cultured according to a previously published protocol, was added to agarose-coated microwells on a 96-well plate (Figure 1A).^36-38^ The hiPSC-VSMC self-aggregated into spheroids within 24 hours without the need for centrifugation. Each microwell consistently formed one round spheroid proportional in size to the number of single cells added. There were four different sizes of spheroids created denoted by the number of cells: 5000, 10000, 20000, and 40000 cells (Figure 1B and 1C). We were also able to make bigger spheroids using 50000, 100000 and 200000 cells (See supplementary Figure 1). Spheroids of each different size had retained their hiPSC-VSMC phenotype as evidenced by their abundant staining for the smooth muscle cell markers calponin, SM-22α, and SMA (Figures 1C and 2). The typical hiPSC-VSMC morphology in a higher magnification can be seen in Figure 2.

**Figure 2:**
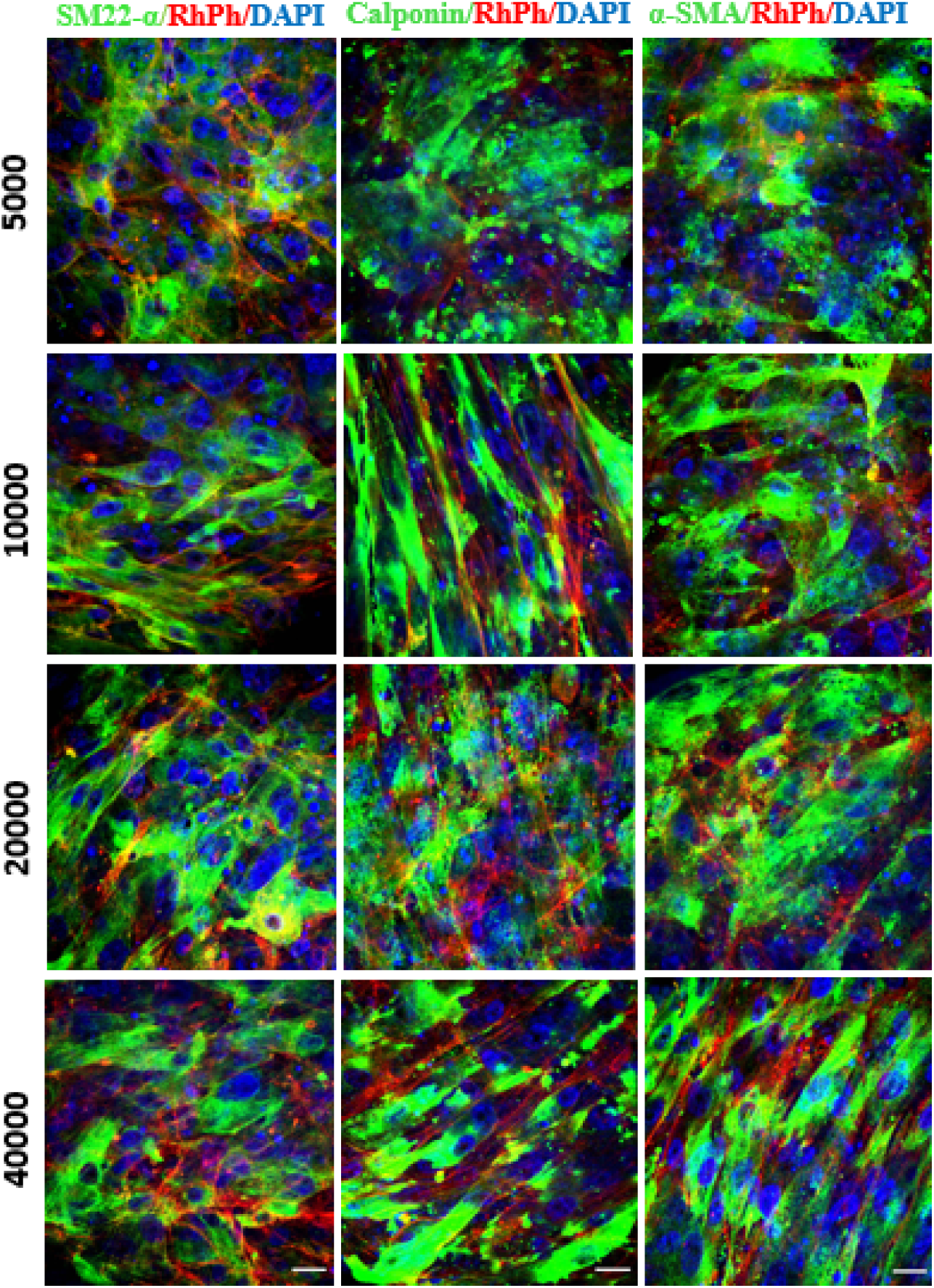
Phenotypic characterization of hiPSC-VSMC. Immunofluorescence images of spheroids composed of 5000 cells, 10000 cells, 20000 cells, and 40000 cells stained with major smooth muscle cell markers SM22-α, Calponin, and SMA in green. The cytoskeleton was stained with rhodamine phalloidin (red), and nuclei with Dapi (blue). Scale bar measures 20 μm.

### Increasing hiPSC-VSMC spheroid size increases cell viability with enhanced LDH level

Relative cell viability within the spheroids was determined by using AlamarBlue assay (Figure 3A). There was a statistically significant increase in cell viability between the 5000-cell spheroid and the 20000-cell and 40000-cell spheroid. The 10000-cell spheroid also had significantly lower viability when compared to the 40000-cell spheroid. However, there was no difference in the relative viability between 5000 and 10000, 10000 and 20000, and 20000 and 40000. The relative viability increased with increasing spheroid size.

**Figure 3:**
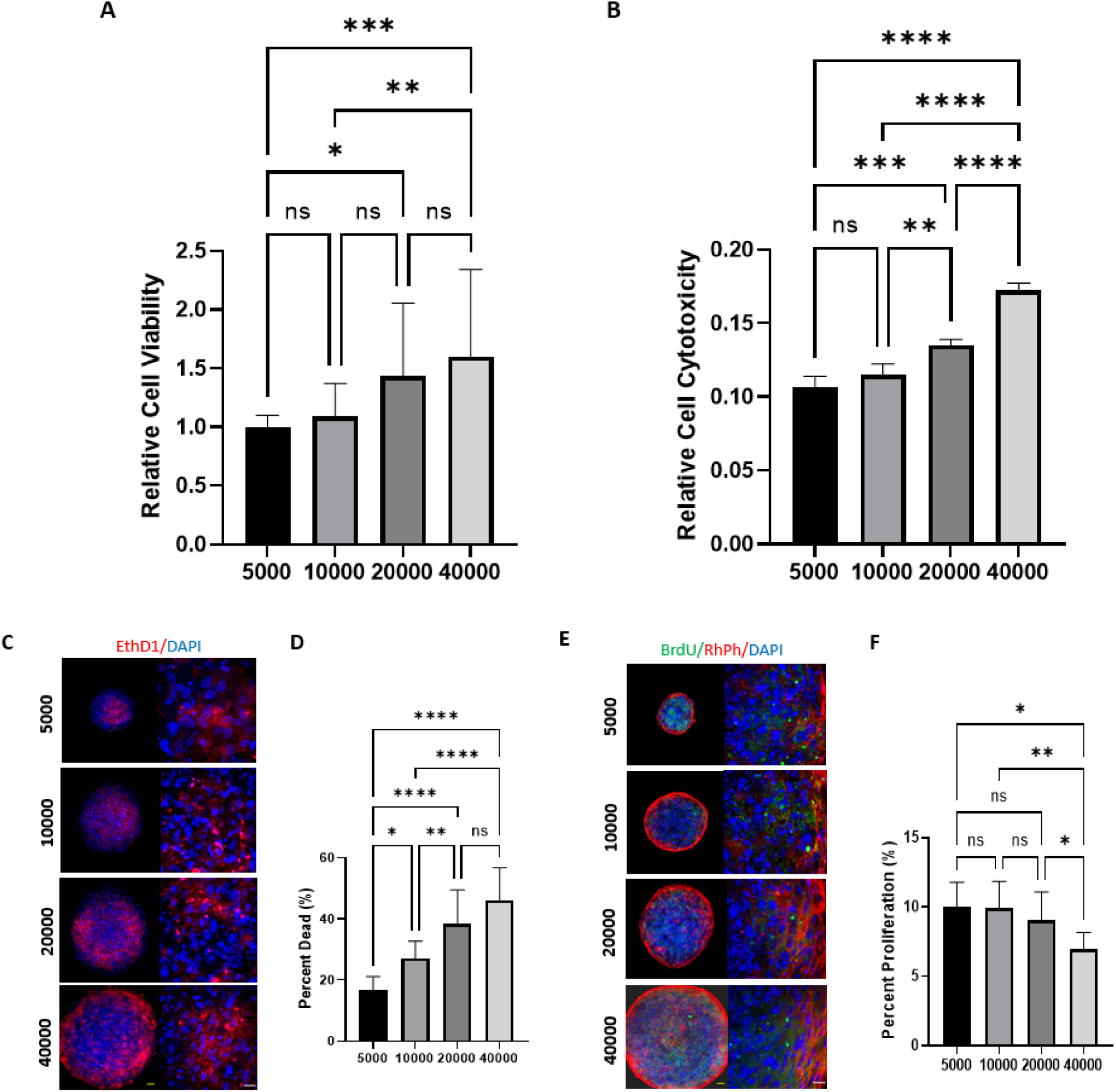
Characterization of the hiPSC-VSMC spheroids for cell viability. Spheroids of various sizes (5000 cells, 10000 cells, 20000 cells, and 40000 cells) were characterized for cell viability. (A) AlamarBlue cell viability assay demonstrating the relative viability of different size spheroids. (B) LDH assay using conditioned medium of different size spheroids to demonstrate cellular cytotoxicity. (C) Representative fluorescence images of live/dead assay comparing relative cell viability of hiPSC-VSMC spheroids. EthD1 stains dead cells (red) and Dapi stains nuclei (blue). Scale bar measures 50 and 20 μm. (D) Quantification of the number of dead cells to total cells presented as a percentage. (E) Representative immunofluorescence images of BrdU proliferation assay of hiPSC-VSMC spheroids. BrdU stains proliferating cells (green), rhodamine phalloidin stains cytoskeleton (red), and Dapi stains nuclei (blue). Scale bar measures 50 and 20 μm. (F) Quantification of proliferating cells based on relative number of BrdU positive cells to total cells and presented as a percentage. * Denotes a statistically significant difference between groups (n = 4–31, one-way ANOVA, *p< 0.05, ***p< 0.001, and ****p< 0.0001)

The relative level of LDH in the conditioned medium was measured to assess the cytotoxicity associated with different sizes of the spheroids mentioned above (Figure 3B). The 40000-cell spheroid had the most amount of LDH while the 5000-cell spheroid had the least. There was no statistically significant difference between the 5000 and 10000-cell spheroids. However, the 20000-cell spheroid had more LDH level than the 10000-cell spheroid. Additionally, the 40000-cell spheroid had more LDH level than both the 10000-cell spheroid and the 20000-cell spheroid. As size increased, so did the level of LDH.

### 10000-cell spheroid is the optimal size with less dead and more proliferating cells

To further characterize the viability of the different spheroids, a cell death assay was done using EthD1 (Figure 3C and 3D). The 5000-cell spheroid had the lowest percentage of dead cells when compared to all the other sizes. The highest percentage of dead cells was seen in the 40000-cell sample. Of note, there was no significant difference in the percentage of dead cells between the 40000-cell spheroid and the 20000-cell spheroid. The 10000-cell spheroid had an intermediate percentage of dead cells with more dead than the 5000-cell sample, but less dead than the 20000 sample.

To determine the number of proliferating cells within the spheroids, a BrdU proliferation assay was carried out (Figure 3E and 3F). The 40000-cell spheroids had the lowest percentage of proliferating cells when compared to all other sizes. However, there was no significant difference in the relative number of proliferating cells between spheroids made of 5000, 10000, or 20000 cells.

### Spheroids bigger than 10000 have no effect on the paracrine secretion of hiPSC-VSMC

Quantitative ELISA was performed to evaluate the relative levels of secreted paracrine factors between the different sizes of spheroids (Figure 4). The 5000-cell spheroids had significantly less relative VEGF secretion than the 10000, 20000, and 40000-cell spheroids, but there was no significant differences between any of the other sizes (Figure 4A). Relative bFGF secretion was the lowest in the 5000-cell spheroids (Figure 4B). There was no significant difference between the levels of bFGF secretion when comparing 10000, 20000, and 40000-cell spheroids. A continuation of this trend was seen with Ang-1, TGF-β, TNF-α, IL-10, SDF-1α, and MMP-2 (Figure 4C – 4H). The 5000-cell spheroids had significantly less secretion of all these factors when compared to their counterparts of a larger size. In figure 4H, the secretion of MMP-2 is seen to be significantly lower in the 10000-cell spheroid compared to the 40000-cell spheroid. Aside from this one notable exception, there were no other significant differences between the levels of factor secretion from the 10000, 20000, and 40000-cell spheroids.

**Figure 4:**
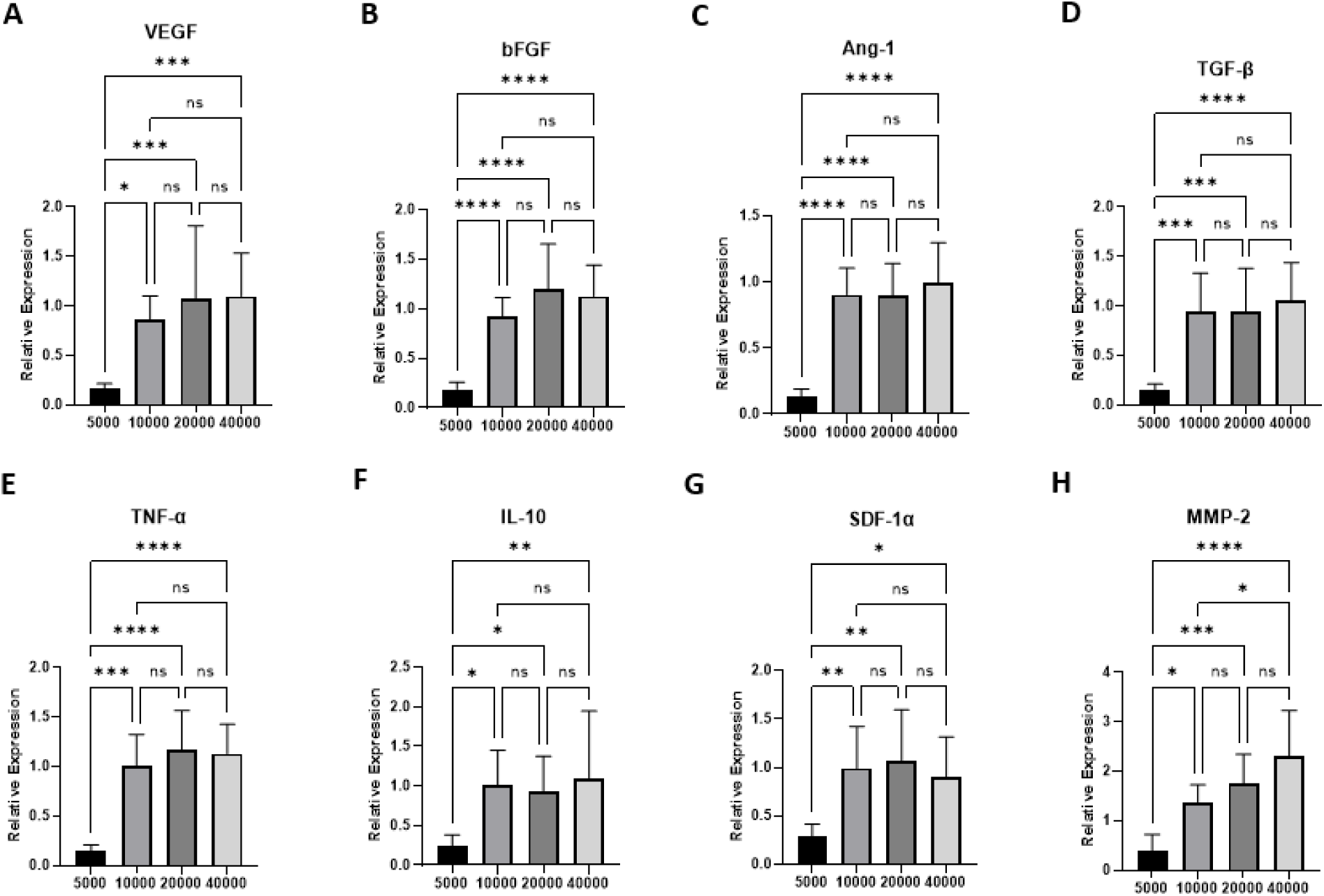
Characterization of paracrine secretion from hiPSC-VSMC spheroids. ELISA was performed using conditioned medium collected from different sized spheroids (5000 cells, 10000 cells, 20000 cells, and 40000 cells). (A) VEGF (*n = 9*) (B) bFGF (*n = 9*) (C) Ang-1 (*n=9*) (D) TGF-β (*n = 8-9*) (E) TNF-α (*n = 6*) (F) IL-10 (*n = 8-9*) (G) SDF-1 α (*n = 8-9*) (H) MMP-2 (*n = 7-9*) * denotes a statistically significant difference between groups (one-way ANOVA, *p< 0.05, ***p< 0.001, and ****p< 0.0001)

### Conditioned medium from spheroids promotes fibroblast proliferation, migration, and paracrine secretion

To characterize the therapeutic benefits of CM from spheroids, a series of functional assays were conducted. The spheroid CM was compared to DMEM, a negative control, and fibroblast CM, a positive control. Primary human skin fibroblasts on a 24-well plate were treated with the three different types of medium. Relative proliferation after treatment was determined using AlamarBlue (Figure 5A). The fibroblasts treated with fibroblast CM and spheroid CM had significantly higher levels of proliferation when compared to the DMEM treatment group. However, there was no difference between cells treated with fibroblast CM and cells treated with spheroid CM. Next, the relative cytotoxicity between the treatment groups was assessed using the LDH cytotoxicity assay (Figure 5B). There was no significant difference in the levels of cytotoxicity between the fibroblasts treated with DMEM, fibroblast CM, and spheroid CM.

**Figure 5:**
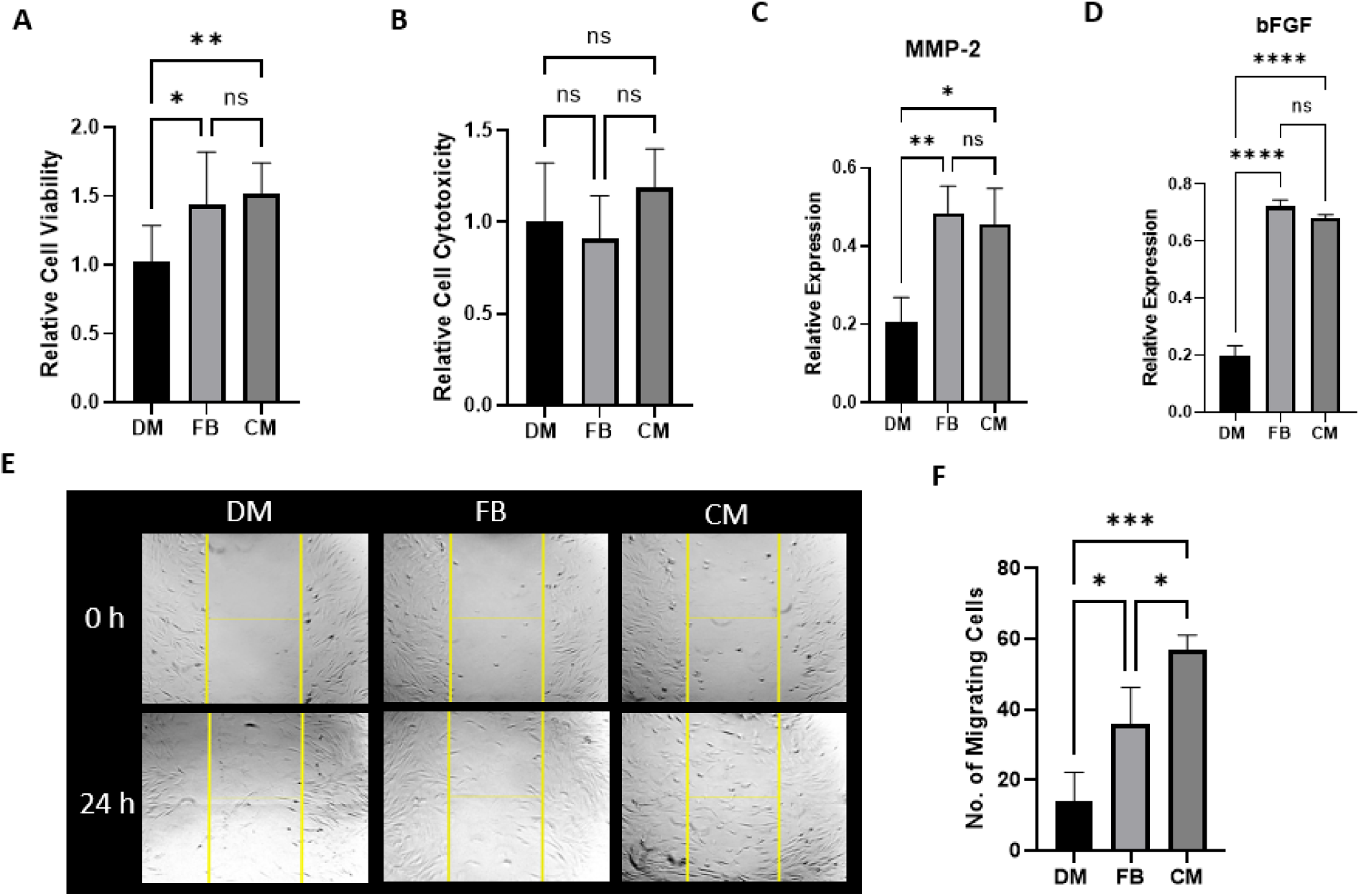
Characterization of conditioned medium from hiPSC-VSMC spheroids. Human dermal fibroblast cells treated with DMEM (DM), fibroblast conditioned medium (FB), and 10000 cell spheroids conditioned medium (CM). (A) AlamarBlue cell viability assay demonstrating the proliferation of different treatment groups. (B) LDH assay of different treatment groups demonstrate relative cellular cytotoxicity. Characterization of paracrine secretion of different treatment groups using ELISA: (C) MMP-2 and (D) bFGF. (E) Bright field images of a migration assay of treatment groups at 0 hours and after 24 hours. (F) Quantification of the migration assay after 24 h. * denotes a statistically significant difference between groups (n = 4 –8, one-way ANOVA, *p< 0.05, ***p< 0.001, and ****p< 0.0001)

To assess how treatment with CM affected the paracrine secretion of fibroblast cells, qualitative ELISA using bFGF and MMP-2 was done. The spheroid CM treated cells expressed higher levels of bFGF when compared to the control, DMEM, as did the fibroblast CM treated cells, but there was no difference in expression between spheroid CM and fibroblast CM. The same results were observed with relative expression of MMP-2. Spheroid CM treated cells expressed higher levels than DMEM and fibroblast CM treated cells also expressed higher levels than DMEM. There was no significant difference between the fibroblast CM and spheroid CM treatment groups.

Migration of fibroblasts post-treatment with DMEM, fibroblast CM, and spheroid CM was assessed via bright field images (Figure 5E) and quantified with manual counting (Figure 5F). The DMEM treatment group had the least migration when compared to the fibroblast CM group and the spheroid CM group. The cells treated with spheroid CM demonstrated significantly enhanced cell migration over the fibroblast CM cells.

## Discussion

Human iPSCs have become central to interest surrounding regenerative cell-based therapies. To realize their full therapeutic potential, there remains a need to optimize culture conditions to improve cell survival, and paracrine secretion.^27^ We and others have used functionalized biomaterials to enhance stem cell survival, and paracrine secretion both in vitro and in vivo.^36-38^ Spheroids with a 3D tissue-like architecture have been shown to improve cell-survival, paracrine secretion, and improved regenerative potential.^4,6,8,14,20,27^

The purpose of this study was to develop a low-cost and reproducible method to form hiPSC-VSMC 3D spheroids of different sizes and evaluate the individual differences between varying sizes to establish the optimal spheroid parameters. We established a self-aggregation method to form spheroids using just agarose-coated, U-bottom microwells.^44^ In agreement with previous studies, our spheroids self-assembled into a secondary spherical structure within 24 hours without the need of any centrifugation. Spheroids of 5000, 10000, 20000, and 40000 cells were consistently, and reproducibly, formed. We also demonstrated that these spheroids maintained their hiPSC-VSMC phenotype as evidenced by their abundant staining with smooth muscle cell markers calponin, SM-22α, and SMA.

Thus far, spheroids have been formed using many different cell types, and at vastly different sizes, with very few studies investigating the effects of spheroid size on important cellular properties. In one of the studies Torizal *et. al*. evaluated the impact of size of the iPSC spheroids on their ability to differentiate into hiPSC-derived hepatocytes. They used 96-well, ultra-low attachment plates and a rotary shaker to form their spheroids and found that increasing size enhanced differentiation. Ultimately, they decided the optimal size for hiPSC-derived hepatocyte spheroids to be 12,500-25,000 cells.^45,46^ In another study Murphy *et. al*. used human bone marrow derived MSCs to elucidate the presence of hypoxia and oxygen tension within the spheroid and the effects oxygen availability on cell survival. They found their spheroids composed of 15000, 30000, and 60000 MSCs had decreasing proliferation with increasing size.^19^ It is clear that there is a diverse array of spheroid types being made with very little consensus on the implications this may have on cellular function.

Although there has been some research using spheroids made with smooth muscle iPSCs, the spheroid size was developed seemingly at random.^47^ We took into consideration the viability, cytotoxicity, proliferation, and paracrine secretion to decide an optimal spheroid size of hiPSC-VSMC for future applications. Our data shows that cellular viability increases with size however we see a cytotoxic effect in the form of LDH release, more dead cells, and less proliferative cells. This is probably due to the fact that cells in 3D culture experience a heterogeneous environment in regards to nutrient supply, oxygen availability, and waste disposal which is ultimately dictated by spheroid size.^22,45,48,49^ As cell number increases, poor oxygenation and nutrient depletion lead to the formation of a necrotic core. ^9,28,45,50^ It has been suggested that the limit of oxygen diffusion *in vivo* is 200 µm with poor conditions found at the core of spheroids larger than 400 µm.^22,28^ In hepatocyte-derived iPSCs, there was no oxygen limitation below 100 µm, but in adipose-derived MSCs, spheroids up to 500 µm were tolerant of hypoxic conditions.^46,49^ Because cells differ in terms of oxygen consumption, metabolic activity, and tolerance for resource depleted environments, studies conducted in cell types other than the cell line of interest may not be as generalizable as we believe.

In the case of hiPSC-VSMC spheroids, this maximal limit may begin somewhere around 40000 cells when cytotoxicity increases and proliferation decreases. Though the 5000-cell spheroid had the lowest cell death and a highest cell proliferation rate, this size did not demonstrate the most growth factor secretion. The remaining sizes all had similar levels of growth factor secretion, but the 10000-cell spheroid had fewer dead cells and more proliferating cells compared to the bigger sizes. We believe the 10000 cell spheroids represent the ideal balance between overall cell health and maximal growth factor secretion for hiPSC-VSMC. For these reasons, we opted to use conditioned medium (CM) from the 10000 cell spheroids for our functional assays.

Cells treated with spheroid CM demonstrated increased proliferation and migration when compared to the control group. Previous studies endorse our findings that CM from cells in 3D culture promote cell mitogenic activity.^11^ The ability to enhance migration and also maximize growth factors, such as bFGF and MMP-2, has promising implications for biological processes, such as wound healing, that are dependent on these mechanisms.

## Conclusion

Our work has demonstrated that spheroid size has an enormous impact on the properties of hiPSC-VSMC including consequences for overall cellular health and paracrine secretion. Future studies should take the optimal spheroid size into consideration. A balance between cell viability and maximizing paracrine secretion must be achieved. However, spheroid size remains tailored to the individual needs and expectations of the researcher. Going forward, more consideration should be given to spheroid size before using them for regenerative healing applications.

## Author Contributions

B.C.D and H.C.H conceived the study and procured the funding. S.I, B.C.D and H.C.H designed the experiments. B.C.D, S.I, and J.P performed the experiments. S.I and B.C.D wrote the manuscript. All the authors participated in data analysis, discussed the results, and reviewed the manuscript.

## Acknowledgements

The authors would like to thank the core research facilities at the Yale Department of Surgery.

