## Supplemental Information for "Human iPSC-Vascular Smooth Muscle Cell Spheroids Demonstrate Size-dependent Alterations in Cellular Viability and Secretory Function"

Running Title: hiPSC-VSMC derived Spheroids for Regenerative Therapy

Address: Yale University School of Medicine, 330 Cedar St., BB 3rd Floor, PO Box 208041, New Haven, Connecticut 06510, Telephone: 203 737-2049, Fax: 203 785-5714

**Materials:**

Vascular smooth muscle cell (SmGM-2) growth media purchased from Promocell, Germany. mTESR medium and dispase from STEMCELL Technologies; 96-well ultra-low attachment and ELISA (NUNC) plates from Thermo Fisher Scientific; matrigel from Corning; agarose from AmericanBio; Rhodamine Phalloidin from Invitogen (Thermo Fisher); TMB substrate from Abcam; and AlamarBlue from Thermo Fisher Scientific. All other reagents and laboratory supplies were purchased from Thermo Fisher Scientific, USA.

**Table-S1:** Information related to primary antibodies.

| Primary Antibody | Dilution | Catalog Number/ Manufacturer |
| --- | --- | --- |
| bFGF | 1:2500 ELISA | 500M38 (PeproTech) |
| VEGF | 1:2500 ELISA | AB-119 (Abcam) |
| Ang-1 | 1:2500 ELISA | MAB9231 (R&D Systems) |
| TGF-β | 1:2500 ELISA | SC-52893 (Santa Cruz Biotechnology) |
| TNF-α | 1:2500 ELISA | SC-52746 (Santa Cruz Biotechnology) |
| IL-10 | 1:2500 ELISA | AB34834 (Abcam) |
| SDF-1 α | 1:2500 ELISA | 500-P87A-100UG (PeproTech) |
| MMP-2 | 1:2500 ELISA | AF902 (R&D Systems) |
| SM-22α | 1:250 IF | AB-10135 (Abcam) |
| SMA | 1:250 IF | SC-32251 (Santa Cruz Biotechnology) |
| Calponin | 1:250 IF | C2687-100UL (Sigma Life Sciences) |
| BrdU | 1:250 IF | SC-32323 (Santa Cruz Biotechnology) |

| Secondary Antibody | Dilution | Catalog Number/ Manufacturer |
| --- | --- | --- |
| Anti-Mouse-HRP | 1:2500 ELISA | AB6789 (Abcam) |
| Anti-Goat | 1:2500 ELISA | AB6789 (Abcam) |
| Anti-Rabbit | 1:2500 ELISA | 31460 (ThermoFisher) |
| Anti-Mouse-Alexafluor488 | 1:250 IF | A21202 (ThermoFisher) |
| Anti-Goat-Alexafluor555 | 1:250 IF | A21432 (ThermoFisher) |

**Table-S2:** Information related to secondary antibodies.

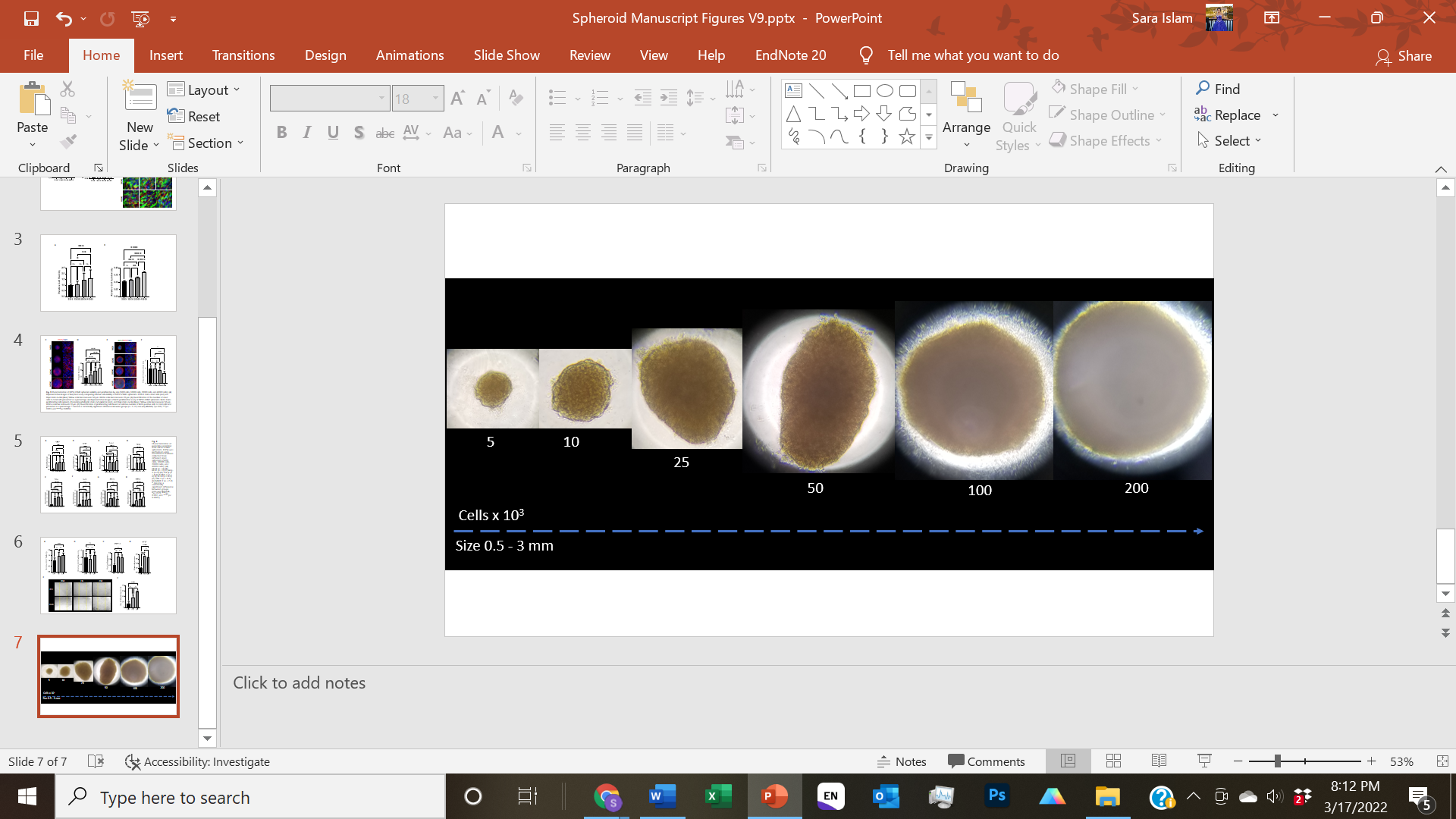

**Supplementary Figure 1:** Spheroid formation. Spheroids formed from varying amounts of cells. Representative bright field images of spheroids. From left-to-right, cell counts are 5000, 10000, 25000, 50000, 100000, and 200000 cells. Scale bar measures 50 μm.
